# A genetic toolkit for stable transgenesis in the anaerobic gut parasite *Blastocystis* ST7-B

**DOI:** 10.64898/2026.04.28.721505

**Authors:** M. Rey Toleco, Kevin S W Tan, Mark van der Giezen

## Abstract

*Blastocystis* is among the most prevalent microbial eukaryotes in the human gut, yet it has remained largely inaccessible to functional genetics. Here, we report a combinatorial toolkit for *Blastocystis* ST7-B that enables stable transgene maintenance under antibiotic selection and recovery of colony-derived transgenic lines. Guided by a proteomics-informed candidate screen, we identified endogenous promoter–terminator pairs and benchmarked their activity using NanoLuc luciferase (Nluc), defining near-background, weak, moderate, and robust expression tiers. We optimised square-wave electroporation and establish conditions that balance DNA delivery with culture viability, providing a practical operating regime for routine transfection. Using resazurin-based viability assays alongside culture outgrowth validation, we identified puromycin and trimethoprim as the most reliable selectable systems. A three-stage workflow combining liquid enrichment, solid-phase selection, and liquid culture expansion supports recovery of colony-derived transgenic lines that can be cryopreserved and revived with retained growth, antibiotic resistance, and reporter expression. Finally, bicistronic constructs incorporating a codon-optimised P2A peptide supported selection-linked expression of anaerobic-compatible reporters (UnaG, smURFP, and SNAP-tag®). Results showed reporter-dependent performance consistent with constraints such as chromophore availability and substrate permeability. Together, this toolkit makes *Blastocystis* ST7-B markedly more amenable to genetic engineering.

## Introduction

*Blastocystis* is an anaerobic microbial eukaryote that colonises the intestinal lumen of its host under extremely low oxygen tension (Stensvold and van der Giezen, 2018). Genetically diverse and morphologically variable, it colonises a wide range of hosts, including humans (Tan, 2008). It is also one of the most prevalent eukaryotic microorganisms in the human gut and is estimated to colonise approximately one to two billion people globally (Scanlan and Stensvold, 2013). Yet mechanistic understanding remains limited, particularly with respect to its metabolic physiology and interaction with its host.

At least 44 distinct subtypes (STs) have been identified based on small subunit ribosomal RNA gene sequences, with ST1 to ST9 found in humans (Koehler et al., 2024; Šejnohová et al., 2024). Among these, ST1 to ST4 are the most frequently detected in human microbiomes (Alfellani et al., 2013). Recently, the presence of *Blastocystis* in the gut was reported to correlate with favourable cardiometabolic profiles and lower BMI values, and to be negatively associated with diseases linked to altered gut ecology (Piperni et al., 2024). However, such generalisations obscure important subtype-specific differences that shape its biology. In animal models, *Blastocystis* ST7-B has been associated with shifts in microbiome composition that include reduced abundance of beneficial taxa, whereas *Blastocystis* ST4-WR1 has been linked to the opposite trend (Yason et al., 2019; Deng and Tan, 2022). In epithelial cell culture, several ST7 isolates trigger tight junction disruption, whereas the ST4 isolates tested did not (Wu et al., 2014). These paradoxical subtype-specific effects complicate any attempt to classify *Blastocystis* as strictly commensal or pathogenic (Deng and Tan, 2025).

*Blastocystis* metabolism is equally unconventional. It encodes both the classical mitochondrial pyruvate dehydrogenase complex and the anaerobic pyruvate:ferredoxin oxidoreductase (PFO), consistent with the potential for dual modes of pyruvate decarboxylation (Stechmann et al., 2008). It encodes an iron-only [FeFe] hydrogenase, typically linked to molecular hydrogen generation, though hydrogen production has not been detected under the conditions tested in *Blastocystis* (Stechmann et al., 2008). Its tricarboxylic acid (TCA) cycle is incomplete, missing key enzymes such as citrate synthase and isocitrate dehydrogenase, and may function in malate dismutation rather than canonical respiration (Stechmann et al., 2008; Gentekaki et al., 2017). Intriguingly, *Blastocystis* expresses alternative oxidase (AOX), a mitochondrial enzyme that uses oxygen as a terminal electron acceptor, although its physiological relevance remains unresolved (Tsaousis et al., 2018). Together, these features reflect the hybrid and evolutionarily distinct nature of its mitochondria.

A further hallmark of *Blastocystis* physiology is mitochondrial glycolysis, where multiple glycolytic enzymes localise to the mitochondria (Abrahamian et al., 2017; Bártulos et al., 2018; Pyrihová et al., 2024). By relocating core carbon flux across the cytosol–mitochondria boundary, this organisation implies rewired redox and ATP economics and makes *Blastocystis* a useful experimental system for testing how metabolism and organelle function co-evolve under anaerobic constraint. A link between mitochondrial glycolysis and serine biosynthesis has been proposed in oomycetes (Abrahamian et al., 2017), but does not apply to Blastocystis, which does not encode mitochondrial versions of the key enzymes required for serine biosynthesis. In parallel, comparative genomics supports a trajectory of reductive evolution, including a streamlined genome and coordinated loss of peroxisomes and flagella, consistent with long-term adaptation to an anaerobic gut niche (Gentekaki et al., 2017; Záhonová et al., 2023).

Though *Blastocystis* occupies a unique position at the intersection of metabolic specialisation, reductive evolution, and global gut colonisation, it remains refractory to molecular-level experimental manipulation. Without functional genetics, mechanistic insight across cellular, metabolic, and ecological dimensions remains speculative. A major step toward genetic manipulation in *Blastocystis* ST7-B was achieved with the development of a protocol for transient transfection (Li et al., 2019). This approach employed a species-specific promoter derived from the upstream region of the legumain gene, paired with its native 3′ UTR containing a conserved motif downstream of the polyadenylation signal essential for expression (Klimeš et al., 2014). Among the regulatory elements tested, this was the only combination shown to reliably drive transgene expression. Furthermore, the system was constrained by low transfection efficiency, dependence on a single validated promoter–terminator pair, and the absence of effective drug-selectable markers at the time. These limitations prevented sustained expression, precluded clonal propagation, and restricted functional studies that rely on persistent gene activity. Addressing these gaps will require further optimisation, including identification of additional endogenous regulatory elements, development of robust selection strategies, and establishment of methods for stable transgene maintenance and clonal line generation. Because transfection has been demonstrated in *Blastocystis* ST7-B, it provides a practical entry point for systematic tool development.

To address these gaps, we developed a genetic toolkit for *Blastocystis* ST7-B. It combines an optimised electroporation workflow for DNA delivery, a screened set of endogenous promoters and terminators for expression control, validated antibiotic markers for selection, and reporter readouts based on P2A-linked constructs and anaerobic-compatible fluorescent proteins.

This combinatorial toolkit establishes *Blastocystis* ST7-B as a genetically accessible system for studying the regulatory logic of its unorthodox cellular physiology and probing organellar evolution under anaerobic constraint. By enabling sustained transgene expression under drug selection, reporter-based assays, and colony-derived transgenic line generation, it provides an entry point for mechanistic experiments on colonisation-relevant traits and subtype-linked host interaction readouts. Beyond its application to Blastocystis, the toolkit offers a transferable framework for molecular tool development in other non-model microbial eukaryotes. By prioritising endogenous regulatory elements, modular construct architecture, and empirical optimisation, this work offers an actionable blueprint for bringing molecular genetics to lineages that remain inaccessible to molecular investigation.

## Methods

### Cell culture

Axenic cultures of *Blastocystis* ST7-B were maintained in pre-reduced Iscove’s Modified Dulbecco’s Medium (IMDM) with stable glutamine in 25 mM HEPES (L0191; Biowest) supplemented with 10% (v/v) heat-inactivated horse serum (Gibco). Cultures were incubated at 37 °C under anaerobic conditions in 2.5 L anaerobic jars with an anaerobic gas pack (Oxoid) for 3 to 4 days, or until a marked increase in cell density was evident, typically accompanied by a colour change in the medium.

### Promoter and terminator selection

Candidate promoter and terminator sources were selected using the omics resources available for *Blastocystis* at the inception of this study. Because no genome-wide promoter map, transcription start site dataset, or experimentally defined regulatory annotation was available for *Blastocystis* ST7-B, we used the published abundance-ranked proteomic dataset for *Blastocystis* ST4-WR1 as a practical starting point for candidate discovery (Armengaud et al., 2017). Specifically, abundant *Blastocystis* ST4-WR1 proteins were used to identify their corresponding annotated genes, after which clear homologous genes were sought in the *Blastocystis* ST7-B genome. We reasoned that ST7-B homologs of highly abundant ST4-WR1 proteins would provide a rational starting set of loci whose endogenous promoter and downstream regulatory regions were more likely to support detectable transgene expression. This rationale was supported by the highly skewed *Blastocystis* ST4-WR1 proteome, in which 193 proteins contribute approximately 50% of the detected proteome and the 13 most abundant proteins account for approximately 10% (Armengaud et al., 2017). This strategy was intended to enrich for candidate regulatory elements for toolkit development, rather than to randomly sample predicted genes or systematically survey the full range of promoter strengths in the *Blastocystis* ST7-B genome.

The 1,000 most abundant *Blastocystis* ST4-WR1 proteins were cross-referenced with the accompanying proteogenomic dataset to identify loci with experimentally supported C-terminal peptide evidence, providing stronger support for the annotated gene-end boundaries used in candidate selection (Armengaud et al., 2017). Corresponding homologs in the *Blastocystis* ST7-B genome annotation, based on Denoeud et al. (2011), were identified using NCBI BLAST searches (Altschul et al., 1990; Camacho et al., 2009) and retained if they showed ≥60% query coverage and ≥75% identity. For each retained *Blastocystis* ST7-B homolog, the putative promoter region was defined as the intergenic sequence upstream of the start codon, extending to the annotated 3′ boundary of the neighbouring upstream gene, as displayed in the NCBI Genome Browser. Where intergenic space was limited, candidate promoter regions sometimes extended into the annotated coding sequence of the neighbouring gene. Alternative promoter lengths were therefore selected according to locus-specific genomic constraints and evaluated individually. Candidate terminator regions were defined operationally as the native 500 bp sequence immediately downstream of the stop codon and were kept constant across promoter-length variants from the same locus to keep the cloning and screening design tractable.

### Cloning

All cloning was performed using the in-house protocol (Toleco and van der Giezen, 2025) adapted from the NEBuilder HiFi DNA Assembly kit (NEB). Plasmids were sent to Microsynth AG (Germany) for Sanger sequencing. For large-scale plasmid preparation, maxi- or giga-preps were carried out using commercial kits (Qiagen) or the Miraprep method (Pronobis et al., 2016). Selection marker genes (Ecdhfr, Hsdhfr, and pac) and anaerobic fluorescent reporter genes (UnaG, smURFP, and SNAP-tag®) were codon-optimized for *Blastocystis* ST7-B and synthesized by Invitrogen GeneArt Gene Synthesis Services (Thermo Fisher Scientific; Supplemental Data 1). All oligonucleotides were designed using Benchling (https://benchling.com) and ordered from Thermo Fisher Scientific (Supplemental Table 1).

The constructs used in this study were derived from the pXS2-P_Legumain_ vector described by Li et al. (2019), which adapted a heterologous expression-vector backbone for transient plasmid-based expression in *Blastocystis* ST7-B. Here, the same molecular backbone was used as a plasmid scaffold carrying Blastocystis-derived regulatory elements and transgenes.

### Estimation of IC_50_

IC_50_ estimation using resazurin-based viability assay was done as described by Mirza et al. (2011). Antibiotics tested were puromycin (Thermo Fisher Scientific), trimethoprim (MedChemExpress), and WR99210 (MedChemExpress); an additional lot of WR99210 was provided by Jacobus Pharmaceutical (NJ, USA) as a generous gift.

### Transfection protocol

The basic transfection protocol was adapted from Li et al. (2019) with modifications. *Blastocystis* ST7-B cells, cultured for 2–3 days, were harvested by centrifugation at 1,000 x g for 5 minutes at room temperature. The pellet was washed with 5 mL of pre-reduced, incomplete cytomix buffer (10 mM K₂HPO₄/KH₂PO₄, pH 7.6, 120 mM KCl, 0.15 mM CaCl₂, 25 mM HEPES, 2 mM EGTA, and 5 mM MgCl₂) and gently vortexed. Cells were pelleted again under the same conditions, and the supernatant was removed by decanting or gentle vacuum suction, leaving approximately 0.5 mL of buffer above the cell pellet. One millilitre of complete cytomix buffer (pre-reduced incomplete cytomix buffer supplemented with 2 mM ATP and 5 mM glutathione, prepared immediately before use) was added, and the pellet was gently resuspended. The resulting approximately 1.5 mL pooled cell suspension was used for total viable cell counting using a hemacytometer.

After counting, the volume corresponding to 5 x 10⁷ cells was transferred to each electroporation reaction and combined with 25 µg of plasmid DNA. The total electroporation volume was adjusted to 500 µL with complete cytomix buffer. After electroporation (GenePulse Xcell™, Bio-Rad), the contents were transferred into a 2 mL microcentrifuge tube using a sterile 1.5 mL Pasteur pipette. The electroporation cuvette was rinsed with 1 mL of complete IMDM medium, and the rinse was pooled with the transfected cells. Cells were centrifuged at 1,000 x g for 5 minutes at room temperature. The supernatant was discarded, and 1.8 mL of complete IMDM was added. The pellet was resuspended by gentle pipetting or flicking of the tube. Further electroporation details are described in the main text. Cultures were incubated under standard conditions inside an anaerobic box (Oxoid) with an anaerobic gas pack (Oxoid), as described above.

### Nluc assay

Promoter–terminator activity was assessed 16–18 h after electroporation using a transient NanoLuc assay adapted from Li et al. (2019), with modifications. This early time point was selected to capture reporter output within the transient-expression window after DNA delivery, before prolonged plasmid loss, differential outgrowth, or culture-level changes could dominate the readout. Because the constructs did not contain a selectable marker, the measured NanoLuc signal reflects early transient reporter output rather than promoter strength independent of DNA uptake, early plasmid retention, or post-transfection recovery. Cells were harvested 16–18 hours post-transfection by centrifugation at 10,000 x g for 2 minutes at 4□°C. The supernatant was discarded, and the pellet was washed with 500□µL sterile PBS. After a second centrifugation under the same conditions, the pellet was resuspended in 350□µL PBS, and 200–300 mg of sterile zirconia beads (Ø 0.1□mm; Biospec) were added. Tubes were placed on ice, and the TissueLyser LT (Qiagen) adapter was flash-frozen in liquid nitrogen for no longer than 2 seconds. Lysis was carried out using the TissueLyser LT at 50□Hz for two cycles of 2 minutes each, with cooling on ice between cycles. The lysate was clarified by centrifugation at 17,000 x g for 10 minutes at 4□°C, and the supernatant was transferred to a fresh 1.5□mL microcentrifuge tube to serve as crude lysate for luciferase assays. For each reaction, 100□µL of crude lysate was mixed with 50□µL Nano-Glo® Luciferase Assay Reagent (Promega) in a luminescence-compatible 96-well plate, protected from direct light, and incubated at room temperature for 10 minutes. Luminescence was measured using a SpectraMax iD5 plate reader (Molecular Devices) following 30 seconds of orbital shaking. All assays were performed in three independent electroporation runs and analysed in technical duplicates; mock-transfected (no plasmid DNA) cells were included as a negative control.

### Selection strategy and clonal propagation

The selection strategy was carried out in phases: liquid phase and solid phase selections followed by re-establishment of the liquid culture. In the liquid phase selection, transfected cells were allowed to recover and propagate until the medium turned yellow without drug treatment, usually within 2 days post-transfection. Cells were then pelleted by centrifugation at 1,500 x g for 5 minutes at room temperature. The supernatant was discarded, and the pellet was resuspended in fresh, pre-reduced IMDM supplemented with antibiotic. The antibiotic concentrations and the graded selection strategy are described in detail in the main text.

Once cells had adapted to high-dose antibiotic concentrations, they were subjected to solid-phase selection. Cultures were diluted with pre-reduced PBS or IMDM to a final density of 1,000 cells/mL. A 200 to 300 µL aliquot of the diluted cell suspension was spread onto pre-reduced IMDM plates containing 1% agar and 0.1% sodium thioglycolate using a sterile cell spreader (Tan et al., 1996a; Tan et al., 1996b; Tan et al., 2000). Plates were incubated overnight, agar-side down, to allow cells to settle. The following day, plates were inverted and incubated for 10–14 days until visible colonies appeared. Individual colonies were picked and transferred to 2 mL microcentrifuge tubes (with perforated lids for gas exchange), containing 1.8 mL of fresh, pre-reduced IMDM and cultured until the medium turned yellow. These clonal lines were subjected to the same liquid phase selection regime and ultimately maintained at a high drug dose referred to as the maintenance concentration.

All incubations were carried out at 37 °C under anaerobic conditions as described above. For contamination control, media were supplemented with an antibiotic cocktail (ABC) composed of penicillin G (100 U/mL), streptomycin (100 µg/mL), levofloxacin (25 µg/mL), and polymyxin B (100 U/mL).

### Confocal Laser Scanning Microscopy

Sample preparation for confocal laser scanning microscopy (CLSM) was adapted from Pyrihová et al. (2024), with minor modifications. All imaging experiments were carried out on circular No. 1.5 glass coverslips (Marienfeld) pre-coated with poly-D-lysine (Gibco).

For UnaG visualisation, washed transgenic *Blastocystis* ST7-B cells were treated with 50 µM bilirubin (BR; Thermo Fisher Scientific) and incubated anaerobically at 37 °C for 1 h. Cells were washed twice with PBS before mounting. Wild-type (WT) cells processed in parallel served as controls.

For smURFP imaging, prepared transgenic cells were permeabilised with 0.1% saponin (Thermo Fisher Scientific) for 10 min and washed twice with pre-reduced PBS. Permeabilised cells were then incubated with 50 µM biliverdin (BV; Merck) under anaerobic conditions at 37 °C for 2 h, washed twice with PBS, and mounted for imaging. WT cells treated identically were used as controls.

For SNAP-tag® imaging, cells were incubated with SNAP-Cell® TMR-Star (NEB) at 0, 3, and 5 µM concentrations for 30 min at 37 °C under anaerobic conditions. Cells were washed twice with PBS prior to mounting. The same SNAP-tag®–expressing *Blastocystis* ST7-B line incubated without substrate was used as a negative control.

All imaging protocols used the same wash regime to remove excess substrate, and cells were counterstained with Hoechst 33342 (Molecular Probes) to visualize DNA. Cells were mounted in ProLong Gold Antifade Mountant (Molecular Probes), sealed with colourless nail polish, and cured overnight prior to image acquisition.

Confocal image stacks were acquired on a Leica TCS SP8 laser-scanning confocal microscope equipped with a 63x/1.40 NA oil-immersion objective (Leica Microsystems). Z-stacks were collected using system-optimized sampling compliant with the Nyquist–Shannon sampling theorem and LAS X LIGHTNING deconvolution-optimized settings (Leica Microsystems), except during SNAP-tag® imaging due to technical issues with our microscope. Laser power, pinhole diameter and other imaging settings were kept constant within each fluorescent reporter system to allow unbiased visual comparison of fluorescence intensities across conditions. For presentation, a subset of optical sections (10 consecutive slices) was collapsed into a maximum-intensity Z-projection. The linear dynamic range for each signal channel was kept constant across treatments.

Single transmitted-light slices were background-corrected using the rolling-ball algorithm (radius 60 px; light background; sliding paraboloid and smoothing enabled) to better visualize cellular morphology relative to the fluorescence signals. The background-corrected transmitted-light images were then overlaid with the corresponding fluorescence channels in the final merged images, and a cell outline was manually traced to approximate cellular boundaries. All image processing was performed in Fiji (ImageJ distribution, version 1.54p).

### Statistical analysis

Data distributions and variance patterns were examined, and the assumptions for parametric testing were not met. Consequently, group differences were evaluated with the Kruskal–Wallis rank-sum test. When omnibus results were significant, post-hoc pairwise comparisons were performed using Dunn’s test with P-values adjusted for multiple comparisons using the Benjamini–Hochberg method. Unless noted otherwise, tests were two-sided with α = 0.05. Dose–response curves were estimated via nonlinear regression of sigmoidal dose–response models. Analyses were conducted in R (version 4.5.1) and GraphPad Prism (10.6.1 (892)).

## Results

### Endogenous promoter–terminator pairs support graded transgene expression in *Blastocystis* ST7-B

Transgene expression in *Blastocystis* ST7-B has so far relied on a single validated endogenous promoter–terminator pair from the legumain locus (LeguP1; Li et al., 2019). To expand the available regulatory parts, we screened 23 NanoLuc reporter constructs containing putative endogenous promoter–terminator pairs from 11 of 14 candidate loci; three loci could not be cloned after two independent attempts and are indicated in Table 1. Each construct paired a candidate upstream promoter region with the corresponding downstream terminator region from the same locus, defined here as the native 500 bp sequence immediately downstream of the stop codon. Where multiple promoter lengths were tested for the same locus, the terminator fragment was kept constant (Table 1; Figure 1A).

**Figure 1.**
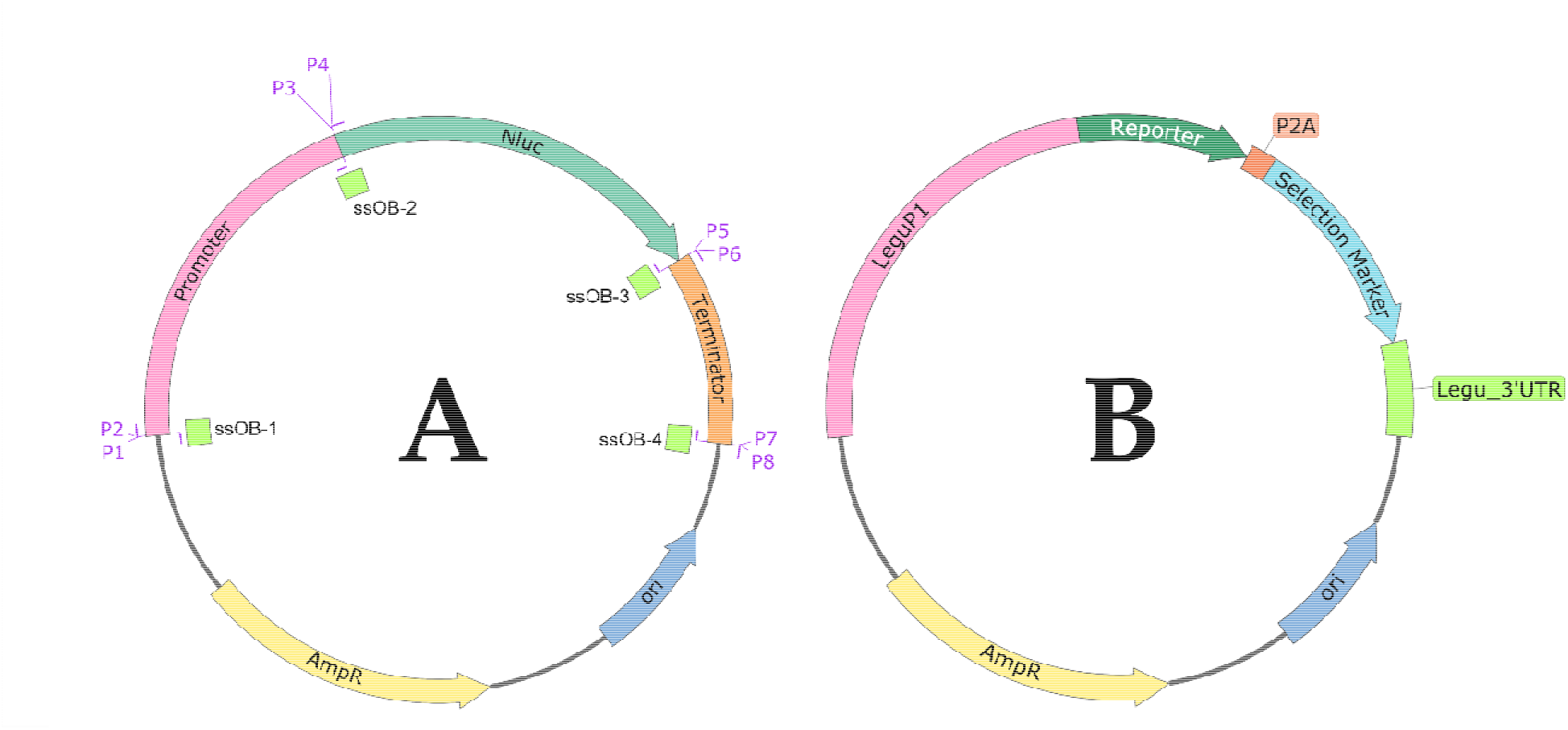
**(A)** Schematic of the plasmid architecture used to assay promoters and terminator using Nluc as a reporter. Coloured blocks/arrows denote interchangeable modules; P1-P8 (purple) represent the necessary primers to PCR amplify the DNA fragments for assembly and green fragments represent the ssOBs needed to complete the DNA assembly reaction. This configuration highlights the modularity of the approach. Plasmids were cloned and amplified in E. coli and then introduced into *Blastocystis* ST7-B, functioning in practice as an E. coli↔*Blastocystis* ST7-B shuttle vector. Robust expression and drug selection were obtained without adding a Blastocystis-specific or canonical eukaryotic origin of replication (e.g., 2µ in yeasts or SV40 ori in mammalian cells), indicating that sequences in the bacterial backbone were sufficient for plasmid maintenance in *Blastocystis* ST7-B. **(B)** A bicistronic construct model using P2A peptide to link fluorescent reporter and a selectable marker under a single promoter-terminato cassette. Reporters used were UnaG, smURFP, and SNAP-tag®; selection markers were Ecdhfr (trimethoprim) and pac (puromycin).

**Table 1.** Genes from which endogenous promoters and terminators were cloned to drive transgene expression in *Blastocystis* ST7-B.

| *Blastocystis* ST4-WR1 |  |  | *Blastocystis* ST7-B |  |  |  |  |
| --- | --- | --- | --- | --- | --- | --- | --- |
| Abundance Ranking* | NCBI protein accession No. | Protein Name† | ST7-B Gene Homolog | NCBI RefSeq | % query coverage | % Identity | Promoters Tested‡ |
| 6 | XP_014525968.1 | aspartate ammonia-lyase; AAL | GSBLH_T000 03689001 | NW_01317 1849.1 | 96.00 | 89.67 | AALpro-1000; AALpro-1500 |
| 8 | XP_014525496.1 | small ADP ribosylation factor GTP-binding protein; SARFGBP | GSBLH_T000 04960001 | NW_01317 1868.1 | 100.00 | 85.39 | SARFGBPpro-600; SARFGBPpro-1325 |
| 9 | XP_014529056.1 | 2-amino-3-ketobutyrate coenzyme A ligase; AKBCoAL | GSBLH_T000 00411001 | NW_01317 1815.1 | 69.00 | 82.98 | Cloning Unsuccessfuls |
| 15 | XP_014525225.1 | 60S ribosomal protein L3; 60SRPL3 | GSBLH_T000 04213001 | NW_01317 1865.1 | 100.00 | 83.22 | 60SRPL3pro-500; 60SRPL3pro-1000 |
| 21 | XP_014529132.1 | aspartate aminotransferase; AAT | GSBLH_T000 01883001 | NW_01317 1821.1 | 89.00 | 85.14 | AATpro-1000; AATpro-1500; AATpro-2438 |
| 32 | XP_014527004.1 | glycine dehydrogenase; GDH | GSBLH_T000 02695001 | NW_01317 1827.1 | 90.00 | 76.32 | GDHpro-1500 |
| 87 | XP_014525499.1 | glyoxysomal and mitochondrial malate dehydrogenases; G/M_MDH | GSBLH_T000 04946001 | NW_01317 1868.1 | 96.00 | 82.59 | G/M_MDHpro-1000; G/M_MDHpro-1500 |
| 124 | XP_014529593.1 | calcium binding mitochondrial carrier protein; CBMCP | GSBLH_T000 03387001 | NW_01317 1838.1 | 100.00 | 81.97 | CBMCPpro-378; CBMCPpro-572 |
| 129 | XP_014528036.1 | 40S ribosomal protein S7; 40SRPS7 | GSBLH_T000 05110001 | NW_01317 1868.1 | 81.00 | 79.93 | 40SRPS7pro-1000; 40SRPS7pro-1500; 40SRPS7pro-2041 |
| 348 | XP_014526323.1 | mitochondrial phosphate transporter; MPT | GSBLH_T000<br>02675001 | NW_01456<br>9325.1 | 100.00 | 80.42 | MPTpro-903;<br>MPTpro-1688 |
| 535 | XP_014529047.1 | cathepsin B1; CathB1 | GSBLH_T000<br>05101001 | XP_0128995<br>54.1 | 96 | 80.32 | Cloning Unsuccessful |
|  |  | cathepsin B2; CathB2 | GSBLH_T000<br>05101001 | XP_0128995<br>55.1 | 90 | 79.84 | CathB2pro-500;<br>CathB2pro-1000 |
| 595 | XP_014525961.1 | EamA-like transporter family;<br>EamA | GSBLH_T000<br>04551001 | NW_01317<br>1866.1 | 97 | 75.51 | EamApro-1023;<br>EamApro-1245 |
| 1000 | XP_014526708.1 | peptidase C1A family protein;<br>PC1AFP | GSBLH_T000<br>01253001 | NW_01317<br>1817.1 | 67 | 77.31 | Cloning Unsuccessful |

\*Based on the *Blastocystis* ST4-WR1 proteome dataset (Armengaud et al., 2017).
†Protein names are from existing annotations in ST4-WR1 or ST7-B; given abbreviations are arbitrarily assigned for nomenclature convenience.
‡Promoter naming is based on the protein abbreviations used in this work followed by the number of bases upstream of the start codon. For example, AALpro-1000 means a genomic sequence of 1000 bp upstream of the start codon of aspartate ammonia-lyase (AAL) in *Blastocystis* ST7-B
§Unsuccessful cloning indicates failure after two independent cloning attempts.

This design allowed construct-level benchmarking of paired promoter–terminator modules, but did not test promoter strength or terminator activity independently. Because the constructs lacked a selectable marker and were assayed 16–18 h after electroporation, NanoLuc activity should be interpreted as early transient reporter output rather than absolute promoter strength. Reporter output across three independent transfections per construct spanned a broad range, from near-background to values approaching the LeguP1 reference, with some individual replicates exceeding the reference measured in parallel. Robust signals were obtained with SARFGBPpro-600, 60SRPL3pro-1000, and EamApro-1023, while 60SRPL3pro-500, EamApro-1245, AATpro-1000, and SARFGBPpro-1325 showed moderate activity (Figure 2A). These constructs therefore expand the set of endogenous regulatory elements that support detectable to strong transgene expression in *Blastocystis* ST7-B.

**Figure 2.**
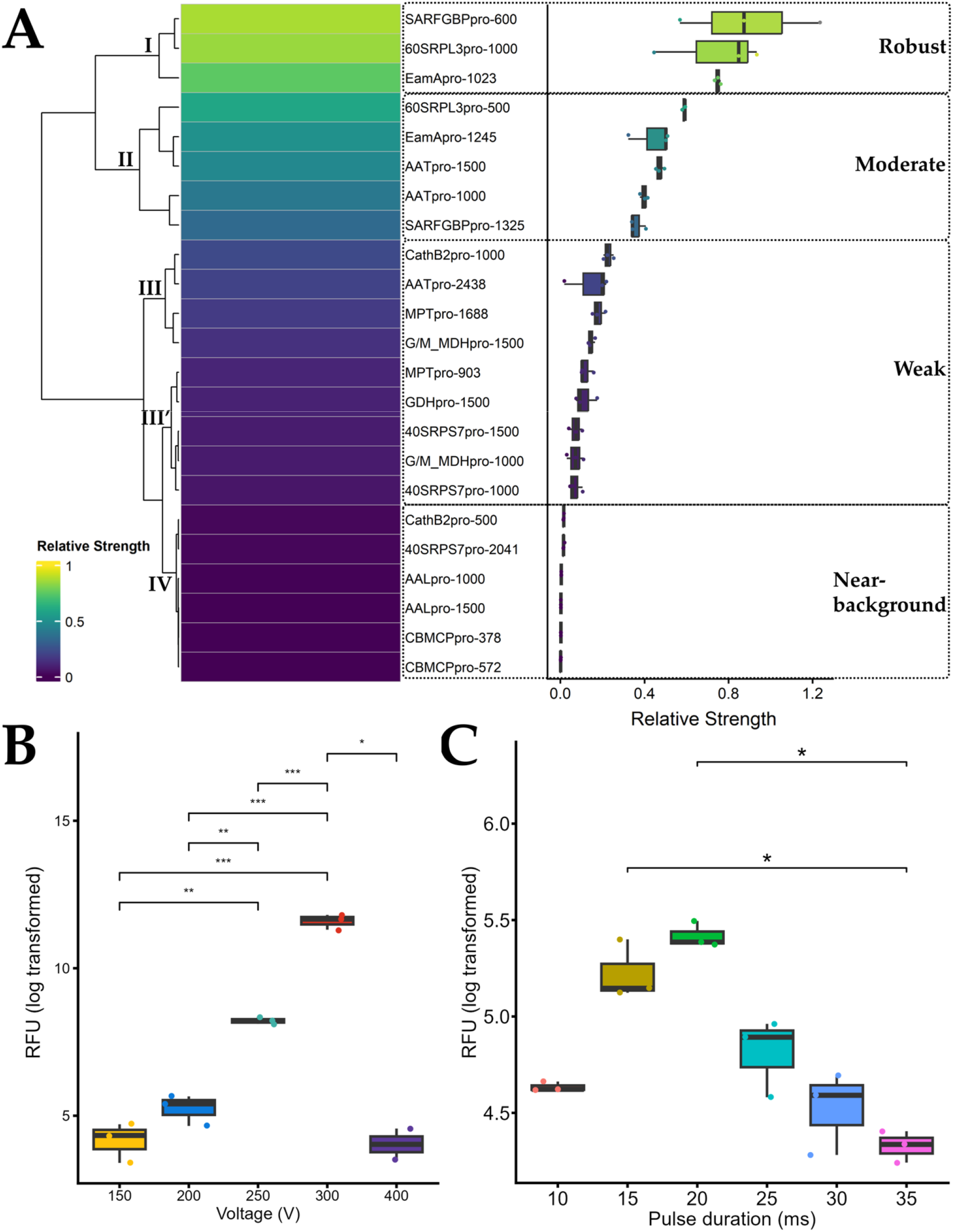
**(A)** Heatmap with hierarchical clustering (left) and boxplots (right) of promoter–terminator activity in *Blastocystis* ST7-B. Activity was measured by Nluc luminescence. For each construct, raw relative light units (RLU; n = 3 independent replicates) were normalized to the LeguP1 reference construct (external standard) to obtain relative strength. Values are displayed on a 0–1 scale using min–max scaling after normalization; on this scale, 1.0 corresponds to the LeguP1 reference. Promoter fragment lengths are indicated by the numbers in the labels. Unsupervised clustering resolved into tiers: (I) robust, (II) moderate, (III and III′) weak, and (IV) near-background. In the adjacent boxplots, points represent values from three independent electroporation runs, each assayed in technical duplicate. **(B)** Effect of square-wave voltage on Nluc luminescence (RLU; n = 3 independent replicates per condition) with pulse duration fixed at 20 ms. **(C)** Effect of single pulse duration on luminescence signal (RLU; n = 3 independent replicates per condition), with voltage fixed at 300 V. For **B** and **C**, electroporation runs were performed with a single pulse while plasmid amount and cell density were held constant. Boxplots show log-transformed raw RLU values (not normalized). Group differences were assessed by Kruskal–Wallis followed by Dunn’s post hoc test (α = 0.05); brackets mark tested contrasts and *, **, *** indicate P < 0.05, 0.01, 0.001, respectively. Although the 20 ms pulse was not significantly different from 15, 25, or 30 ms, it was chosen for subsequent electroporation runs because it produced the narrowest interquartile range (least dispersion) while maintaining a high signal.

Most of the remaining constructs produced weak reporter output relative to LeguP1, and several remained near-background, including AALpro-1000, AALpro-1500, CBMCPpro-378, and CBMCPpro-572. For AAL and CBMCP, increasing promoter length did not improve activity under the conditions tested. Candidate selection had been guided by *Blastocystis* ST4-WR1 protein abundance and the presence of experimentally supported C-terminal peptide, but these features did not reliably predict regulatory performance in *Blastocystis* ST7-B. For example, although AAL is highly abundant in *Blastocystis* ST4-WR1, both AAL promoter fragments drove only near-background expression in *Blastocystis* ST7-B (Figure 2A).

### P2A supports selection-linked co-expression in *Blastocystis* ST7-B

To add a bicistronic expression option in *Blastocystis* ST7-B, we inserted a codon-optimised P2A sequence into a LeguP1-driven cassette linking a selectable marker and UnaG reporter (Figure 1B). The P2A peptide is expected to promote ribosomal skipping during translation, allowing two separate polypeptides to be produced from a single open reading frame. UnaG was used because GFP-family fluorophores require oxygen for chromophore maturation and are therefore unsuitable for anaerobic systems. Before drug selection, fluorescence microscopy of bulk-transfected cultures detected a subset of UnaG-positive cells, indicating expression from the bicistronic construct (Figure 3A). After puromycin selection, UnaG-positive cells were retained and enriched in the surviving population. These results are consistent with functional P2A-linked co-expression of both a selectable marker and a reporter protein in *Blastocystis* ST7-B. However, protein-level separation was not directly tested, and the extent of any residual uncleaved fusion product remains unresolved.

**Figure 3.**
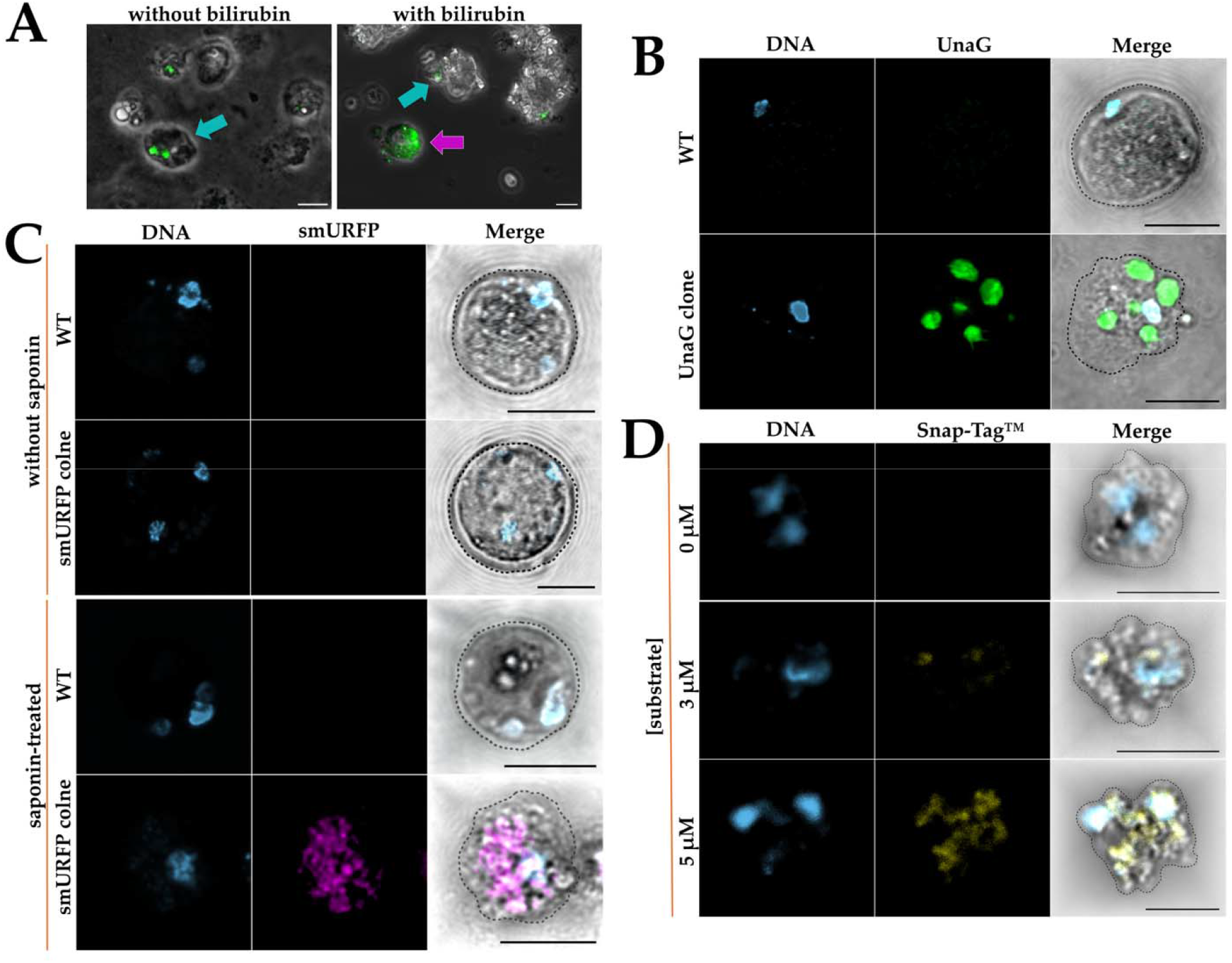
(**A**) Fluorescence micrographs of *Blastocystis* ST7-B transfected with LeguP1:UnaG::P2A::pac:LeguTer and imaged 3 days post-transfection before antibioti selection using an Olympus IX81 fluorescence microscope. Images show phase-contrast overlaid with the UnaG channel. In the absence of exogenous BR, only weak signal i detected, consistent with reported intrinsic autofluorescence in *Blastocystis* (cyan arrows; Nagel et al., 2015). After incubation with 1 mM BR, UnaG-positive cells show strong fluorescence (magenta arrow), whereas neighbouring non-transformed cell remain at background. Scale bar, 10 µm. (B) Representative single-cell confocal images showing compartmentalised UnaG fluorescence in *Blastocystis* ST7-B after incubation with 50 µM unconjugated BR. WT and UnaG-expressing cells were imaged under identical acquisition settings. For display purposes, the lower threshold of the UnaG channel in the transgenic clone was set to the maximum intensity observed in the WT control, so that only signal above the WT autofluorescence range is visible. (C) smURFP fluorescence in *Blastocystis* ST7-B is limited by chromophore access and is unmasked by saponin permeabilisation. Live cells expressing cytosolic smURFP were incubated with 50 µM BV and imaged by confocal microscopy either without permeabilisation or after treatment with 0.1% saponin. No signal was detected above background without saponin. After permeabilisation, smURFP fluorescence became strong and broadly distributed throughout the cytoplasmic compartment, consistent with the untargeted construct design. (D) SNAP-tag® labelling in *Blastocystis* ST7-B is dependent on substrate availability and concentration and yields broadly distributed fluorescence. Live clones expressing an untargeted cytosolic SNAP-tag® were incubated with increasing concentrations of a cell-permeant benzylguanine fluorophore substrate and imaged by confocal microscopy. Signal was at or near background at low substrate but increased progressively with substrate concentration and became broadly distributed throughout the cytoplasmic compartment, consistent with efficient covalent labelling in live cells. DNA was visualised using Hoechst 33342. Most cells contained two nuclei, and smaller Hoechst 33342-positive signals consistent with mitochondrial DNA were also observed in some instances. Because images for the different reporter systems were acquired under different imaging conditions, they are presented to demonstrate reporter detectability and labelling pattern and should not be used for quantitative comparison of cell morphology across reporter systems. Scale bars in B, C, and D represent 5 µm.

### Square-wave electroporation improves DNA delivery and culture recovery in *Blastocystis* ST7-B

Applying the time-constant electroporation protocol of Li et al. (2019) at the recommended 370 V frequently caused arcing and loss of viability in *Blastocystis* ST7-B. Under these conditions, cells darkened, settled at the bottom of the tube, failed to recover, and cultures showed no colour change after 3 to 5 days of incubation. Lowering the voltage to 350 V eliminated arcing, but transfection outcomes remained inconsistent across replicates, indicating that reducing voltage alone did not resolve the limited reproducibility. We therefore replaced time-constant operation, which produces exponential voltage decay, with square-wave electroporation, which maintains constant voltage, and optimised the settings empirically using the LeguP1-Nluc reporter construct.

Voltage was first tested across 150 to 400 V using a single 20 ms pulse in 4 mm cuvettes, while cell input, DNA amount, and transfection volume were held constant at 5 x 10^7^ cells, 25 µg DNA, and 500 µL, respectively (Figure 2B). At 150 to 200 V, corresponding to 0.375 to 0.50 kV/cm, Nluc signals were low. Reporter output increased sharply at 250 V and peaked at 300 V, corresponding to 0.625 and 0.75 kV/cm, respectively, whereas signal dropped markedly at 400 V even in replicates without visible arcing. Both 250 and 300 V produced strong reporter output without arcing or obvious loss of viability, and cultures showed a colour change within 24 hours, with 300 V giving the strongest performance under matched conditions.

Pulse duration was then tested at fixed voltage by comparing single pulses from 10 to 35 ms at 300 V while holding all other parameters constant (Figure 2C). Reporter output increased from 10 to 20 ms and declined at longer pulse durations, with 20 ms consistently outperforming both shorter and longer pulses across replicates.

Because cuvette geometry, electroporation buffer, DNA amount, cell number, and reaction volume were unchanged across these experiments, the observed differences in reporter output and recovery were most consistent with effects of waveform, voltage, and pulse duration. Square-wave electroporation at 300 V for 20 ms in 4 mm cuvettes was therefore used for subsequent transfections.

### Drug sensitivity profiling identifies candidate selection agents for *Blastocystis* ST7-B

To link transgene carriage to cell survival and enable recovery of stable *Blastocystis* ST7-B transformants, we evaluated candidate drugs to establish practical drug–marker pairs for post-transfection selection. Puromycin, trimethoprim, and WR99210 were prioritised because each has a well-established resistance cassette suitable for vector-based expression, namely pac, E. coli dhfr (Ecdhfr), and human DHFR (Hsdhfr), respectively (Vara et al., 1985; Asselbergs and Widmer, 1995; Fidock and Wellems, 1997; Remcho et al., 2020). WR99210 was additionally included because a putative DHFR–thymidylate synthase homolog is encoded in the *Blastocystis* ST7-B genome, providing a rationale to test whether this compound inhibits ST7-B growth.

Dose–response assays were performed using a resazurin-based viability readout to quantify drug sensitivity and define working concentration ranges (Figure 4). Cells were exposed to concentrations ranging from 0 to 1.5 µM for each compound, yielding IC₅₀ values of 30 ± 6.5 nM for puromycin, 102 ± 24 nM for trimethoprim, and 27 ± 13 nM for WR99210. These values confirm that each compound reduces *Blastocystis* ST7-B viability under the conditions tested. These inhibitory profiles identified puromycin, trimethoprim, and WR99210 as candidate selection agents for further testing, with subsequent experiments supporting puromycin and trimethoprim as the most reliable systems under the conditions used here.

**Figure 4.**
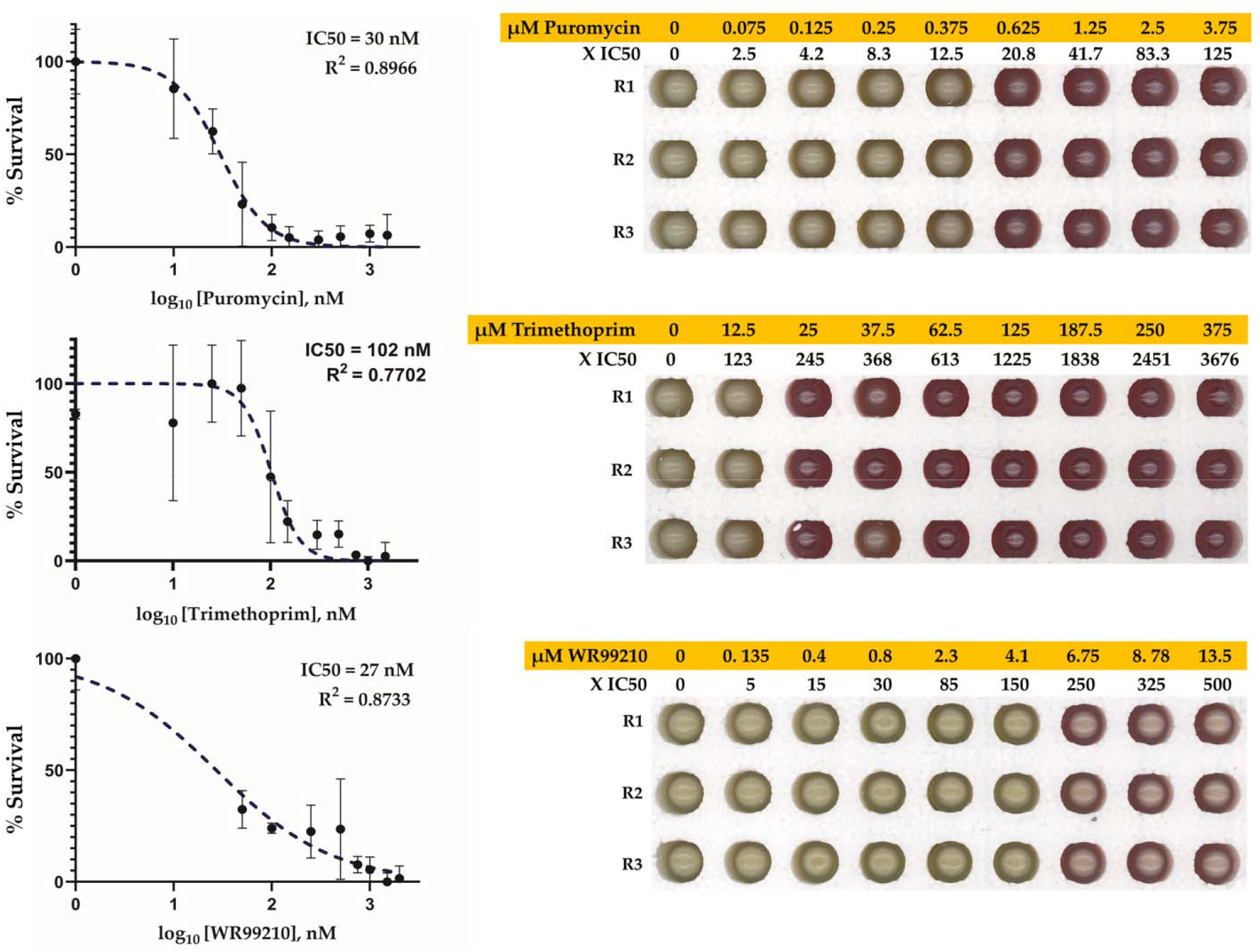
Antibiotic potency and selection-window determination in *Blastocystis* ST7-B. Dose–response curves for puromycin, trimethoprim, and WR99210 were estimated from a resazurin-based viability assay (n = 3 independent replicates per drug pe concentration). Points show mean ± SD, and the insets list the estimated IC_50_ values with R²-values > 0.75 for all fitted curves. The IC₅₀ estimates represent the drug concentrations that reduced resazurin-based metabolic activity by 50% under the assay conditions. Right panels: small-culture complete growth inhibition assay using 1 x 10⁷ WT *Blastocystis* ST7-B cells per culture, assayed in triplicate across a wide range of concentrations. Cultures were incubated for 2 days, and outgrowth was assessed using phenol red acidification of the medium as a culture-level readout, with yellow indicating growth and red indicating no detectable growth. The yellow-to-red transition was used to estimate the concentration required for complete growth inhibition and to guide the subsequent antibiotic selection strategy for *Blastocystis* ST7-B transformants. IC₅₀ and CGI represent distinct assay endpoints: the former measures partial reduction in metabolic activity, whereas the latter identifies the concentration at which no detectable culture outgrowth occurs under the small-culture assay conditions.

### Recovery of colony-derived transgenic *Blastocystis* ST7-B lines through a staged selection regime

To recover *Blastocystis* ST7-B transformants after plasmid transfection, we developed a three-stage workflow comprising stepwise antibiotic selection in liquid culture, isolation of single colonies on solid medium, and re-establishment of individual colonies in liquid culture for expansion and verification. Complete growth inhibition (CGI) was defined as the lowest concentration at which no detectable growth occurred over 24 to 48 h of incubation. Growth and viability were monitored using simple culture phenotypes: healthy cultures shifted the medium towards yellow and formed dense pellets that resisted resuspension after addition of fresh medium, whereas non-viable cultures retained reddish pink medium and produced pellets that dispersed readily. In 1 x 10⁷ WT cells, CGI occurred at 20.8x, 245x, and 250x the provisional IC₅₀ for puromycin, trimethoprim, and WR99210, respectively (Figure 4). Direct transfer of transfected populations into CGI-level drug concentrations led to loss of viability, so selection pressure was increased gradually over several passages. Post-electroporation recovery and selection were performed in 2 mL round-bottom microcentrifuge tubes to conserve space and maintain high cell density. To support anaerobiosis while limiting contamination, each snap-cap lid was perforated with a single pinhole, and drug concentration was increased stepwise by pelleting cells, removing supernatant, and resuspending in fresh medium at the next concentration (Figure 5). Because axenisation is difficult in *Blastocystis* (Chen et al., 1997), recovery media were supplemented with penicillin, streptomycin, levofloxacin, and polymyxin B, which suppressed bacterial contaminants without obvious effects on *Blastocystis* ST7-B growth or morphology during routine monitoring.

**Figure 5.**
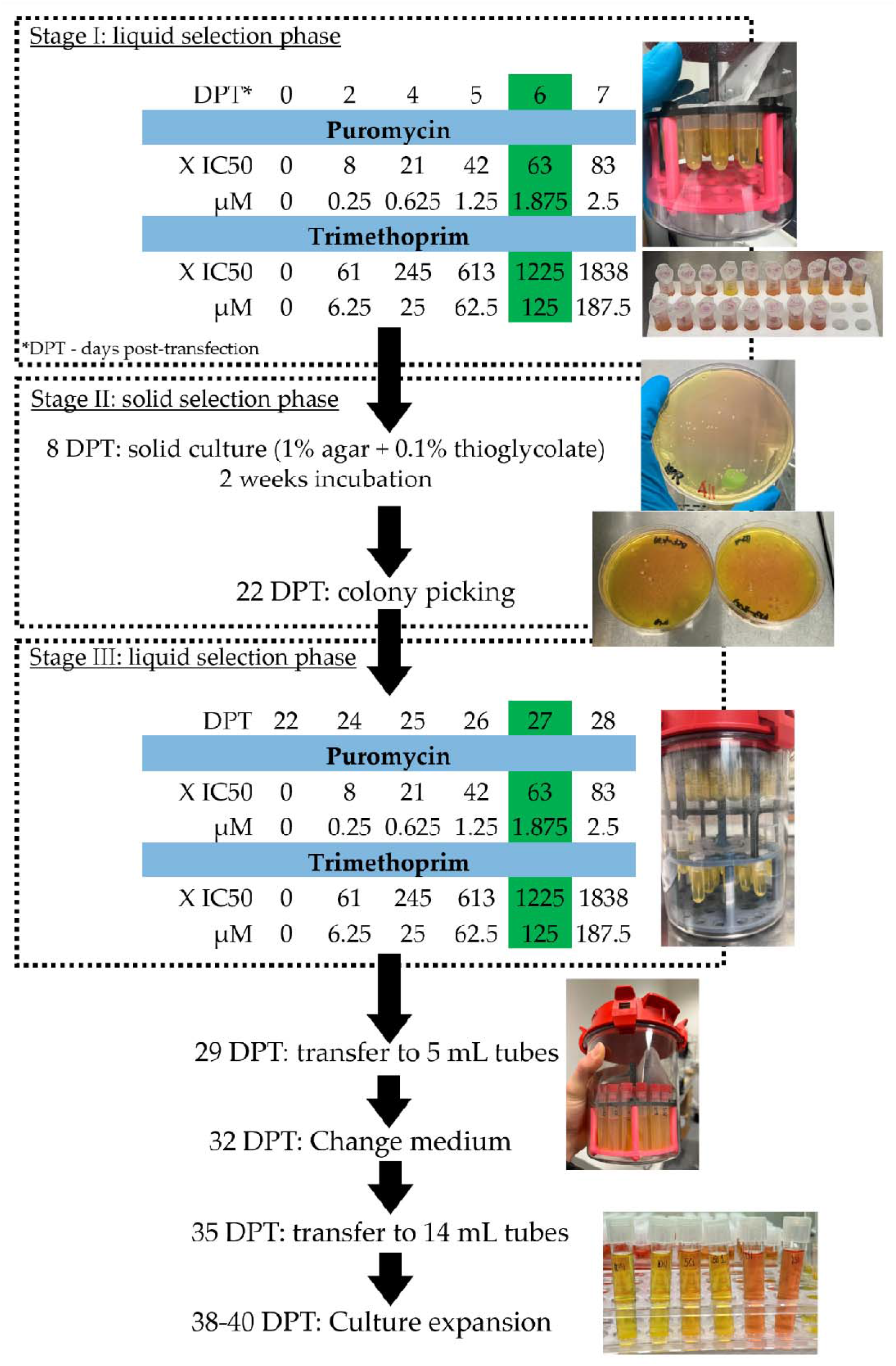
Stepwise selection and clonal propagation of *Blastocystis* ST7-B transformants with puromycin and trimethoprim monitored over days post-transfection (DPT). **Stage I:** transfected cells are kept in 2 mL tubes inside anaerobic jars and exposed to increasing drug concentrations; values highlighted in green refer to the maintenance concentration. **Stage II:** enriched cultures are plated on solid medium incubated for 2 weeks, yielding well-isolated colonies that are picked at 22 DPT. At this stage, the drug concentrations are lowered to ensure that individual plated cells, which later form colonies, remain within their resistance capacity during early outgrowth on the plate. **Stage III:** colonies are re-established in liquid culture under the same stepwise antibiotic regime. Subsequently, volume expansion from 2 mL microcentrifuge tubes to 14 mL culture tubes proceeds through an intermediate 5 mL step to increase the success of culture establishment. The entire workflow takes up to 40 days.

The solid-phase step yielded colony-derived lines consistent with outgrowth from individual viable units, although not all colonies resumed growth after transfer back into liquid culture, indicating strong density dependence during outgrowth. Recovery improved when plates were inoculated at low density, approximately 200 cells per plate, and incubated for up to 14 days before colony picking. Verification at this stage relied on phenotype and sustained growth under the specified sub-CGI drug concentration, as no growth was observed on plates supplemented at CGI levels, and mock-transformed controls plated in parallel also showed no growth. Across passages, puromycin and trimethoprim consistently discriminated transformed from non-transformed populations. By contrast, WR99210 showed marked source-dependent variability: an initial lot supported inhibition and selection at the expected levels, whereas an independently sourced lot, including freshly prepared stocks, failed to inhibit growth even at concentrations up to 1500× the earlier IC₅₀ estimate, and low-passage WT controls no longer reproduced the previous response pattern. ¹H and ¹³C NMR spectra provided no evidence of WR99210 degradation (data not shown). WR99210 was therefore deprioritised, and puromycin and trimethoprim were used for routine selection in *Blastocystis* ST7-B under these conditions. To reduce repeated sampling of the same lineage during recovery and expansion, three to five electroporation runs were performed per construct, and at least one robust clone from each run was advanced. This workflow enabled recovery of colony-derived transformants that retained drug-resistant growth during extended passaging (>15) and after one year of cryogenic storage.

### Anaerobic-compatible fluorescent proteins function in transgenic *Blastocystis* ST7-B lines

To functionally assess transgene expression in *Blastocystis* ST7-B and evaluate the suitability of anaerobic-compatible fluorescent reporters in this organism, three reporters, UnaG, smURFP, and SNAP-tag®, were expressed from bicistronic P2A vectors and examined by confocal microscopy (Kumagai et al., 2013; Rodriguez et al., 2016; Keppler et al., 2003). For each of these proteins, the addition of an external substrate is necessary to form the fluorescent holoenzyme. UnaG, smURFP, and SNAP-tag® require BR, BV, and SNAP-tag® substrates, respectively. Ultimately, each reporter places different demands on chromophore chemistry and substrate permeability. The P2A constructs combined codon-optimized reporters with selectable markers for puromycin or trimethoprim, and stable drug-resistant lines were isolated and expanded for functional characterization of transgene expression.

UnaG, a bilirubin-binding fluorescent protein originally isolated from the muscle of the Japanese eel (Kumagai et al., 2013), produced a bright green signal that rose clearly above WT autofluorescence when background was subtracted under identical settings. Under these conditions, the UnaG signal was readily detectable and visually robust. Efficient holoenzyme formation was observed without permeabilisation of the cells, which indicates that the reporter can become fluorescent in vivo (Figure 3B). The UnaG-derived fluorescence did not appear as a uniform cytosolic signal. Instead, it resolved into discrete, bright subcellular foci and larger compartment-like regions, with darker intervening areas that showed little or no fluorescence. The UnaG coding sequence used in these constructs lacked any intentional targeting peptide or signal sequence, so the underlying plasmid design was expected to drive broadly ectopic expression throughout the cell.

smURFP (small ultra-red fluorescent protein) expression produced a clean far-red signal in *Blastocystis* ST7-B, but only when cells were permeabilised with 0.1% saponin prior to BV treatment (Figure 3C). Under identical laser power and detector settings, non-permeabilised smURFP-expressing cells remained at background levels that were indistinguishable from WT controls. After saponin treatment, smURFP fluorescence rose clearly above autofluorescence and was readily detectable, confirming that the reporter is expressed and can form a functional holoenzyme in *Blastocystis* ST7-B once the chromophore gains access to the cytoplasm. The smURFP signal appeared specific and free of obvious bleed-through into other imaging channels. Within permeabilised cells, the far-red signal was relatively global across the cytoplasm, without strong restriction to discrete subcellular compartments, consistent with the expected distribution of a broadly expressed cytosolic reporter. This pattern matches the construct design, which did not include any intentional targeting peptide or signal sequence and was therefore anticipated to drive ectopic, global expression.

SNAP-tag® labelling was compatible with *Blastocystis* ST7-B and produced robust fluorescence. In cells expressing SNAP-tag®, addition of a cell-permeant SNAP substrate yielded a clear fluorescent signal across a range of substrate concentrations (Figure 3D). Signal intensity scaled with substrate dose over the tested range, indicating that labelling efficiency is substrate-limited under standard conditions. Fluorescence appeared broadly distributed throughout the cytoplasm rather than restricted to discrete organelles, which agrees with the construct design that lacks any explicit targeting sequence and is expected to drive ectopic, global expression. Efficient labelling was obtained without semi-permeabilisation, indicating that the substrate permeated the cell.

## Discussion

This study establishes an integrated genetic toolkit for *Blastocystis* ST7-B that links construct design, DNA delivery, antibiotic selection, clonal recovery, and reporter validation into a single experimental pipeline. Its main advance lies not in any one component alone, but in showing that these elements function together to enable stable transgenesis under antibiotic selection and downstream cell biological analysis in this organism. The toolkit was intentionally designed as a combinatorial system of interchangeable genetic parts, allowing constructs to be tailored to the desired circuitry through promoter choice, P2A-based bicistronic design, and selection markers or oxygen-independent reporters. The previously reported ssOB cloning approach (Toleco and van der Giezen, 2025) further supports this design by enabling straightforward assembly and exchange of parts, giving the system a quasi-modular cloning property.

A key outcome of this study is the expansion of endogenous regulatory elements beyond the legumain module previously used for transgene expression in *Blastocystis* ST7-B (Li et al., 2019). This matters because a useful toolkit requires more than a single high-output cassette. Strong expression may be advantageous for selectable markers and bright reporters, but not for every cargo. Hence, weak or near-background promoter-terminator pairs may be useful for expressing toxic proteins or tightly regulated genes that would otherwise harm the host at high expression levels. The promoter-terminator combinations identified here therefore provide a practical expression range for future reporter design and multi-gene constructs. At the same time, the screen highlights an important constraint on part discovery across *Blastocystis* subtypes. Candidate nomination based on *Blastocystis* ST4-WR1 protein abundance and the presence of experimentally supported C-terminal peptide provided a reasonable starting point, but these features were not sufficient to predict activity in *Blastocystis* ST7-B, as illustrated by the poor performance of the AAL-derived fragments. Cross-subtype information can therefore guide candidate selection but cannot substitute for direct validation in the subtype being engineered. The promoter length series further indicates that regulatory output in *Blastocystis* ST7-B is strongly context-dependent. Longer fragments did not consistently improve activity, arguing against simple length-based design rules and instead pointing to a more compact promoter organisation in which relatively small sequence differences can alter initiation efficiency or introduce inhibitory context (Li et al., 2019). Although the present constructs were not designed to define the regulatory architecture of these promoter regions directly, they point to a clear next step in dissecting this logic, including transcription start site mapping and refinement of promoter boundaries. A similar principle likely applies to 3′ regions, where locus-specific cleavage and yet unknown features may also contribute to the observed differences in expression.

A second important contribution of this study is the refinement of DNA delivery and recovery conditions for *Blastocystis* ST7-B. Electroporation in *Blastocystis* ST7-B appears to operate within a narrow window in which efficient DNA delivery must be balanced against cell survival and post-transfection recovery. Under the previously recommended time-constant settings at 370 V, arcing was frequent and cultures often failed during recovery. By contrast, the optimised square-wave settings avoided arcing and preserved enough viable cells to support post-transfection outgrowth. This distinction matters because successful transfection in *Blastocystis* ST7-B is not defined solely by DNA entry, but also by whether a sufficient number of cells survive to re-establish growth, a practical constraint consistent with the long-recognised but still poorly understood density-dependent growth behaviour of *Blastocystis* in liquid culture (Ho et al., 1993). Selection and clonal recovery then complete the transition from transient uptake to stable antibiotic-selected line generation, and the present data indicate that this recovery phase is a major bottleneck. Immediate transfer into fully inhibitory drug concentrations caused culture collapse, whereas staged enrichment allowed resistant populations to emerge, indicating that stable antibiotic-selected line generation depends as much on managing post-transfection stress as on drug potency itself. Among the selectable systems tested, puromycin and trimethoprim proved the most reliable in practice. WR99210 was initially attractive because of its established use in other protists and the presence of a putative DHFR-TS homologue in the *Blastocystis* ST7-B genome (Fidock and Wellems, 1997; Remcho et al., 2020), but its inconsistent behaviour across sources argues against routine use in Blastocystis. This result is informative because selectable markers that perform well in one protist cannot be assumed to transfer cleanly to another, and in toolkit development practical robustness matters more than theoretical compatibility. This vulnerable recovery phase also places a premium on careful culture handling. Bacterial contamination can readily compromise recovering *Blastocystis* cultures, making contamination control a practical part of workflow feasibility rather than a minor technical detail.

The P2A bicistronic design adds useful flexibility to the system by enabling two coding sequences to be expressed from a single transcriptional unit, for example a selectable marker paired with a reporter. In a platform where fully validated regulatory parts remain limited, P2A offers a practical way to simplify construct architecture while expanding design options. P2A was selected because it is a well-characterised peptide with high reported separation efficiency in human cell lines, zebrafish embryos, and mice (Kim et al., 2011). It also has precedent across microbial eukaryotes, including the protist Dictyostelium discoideum (Zhu et al., 2023), the fungi Aspergillus niger (Schuetze and Meyer, 2017) and Ustilago maydis (Müntjes et al., 2020), and the apicomplexan parasites Toxoplasma gondii (Markus et al., 2019) and Plasmodium falciparum (Dans et al., 2024). However, P2A performance is context-dependent, and the evidence presented here is functional rather than biochemical. P2A-containing constructs support antibiotic-selected recovery and downstream reporter expression in *Blastocystis* ST7-B, but ribosomal skipping efficiency and any residual uncleaved product will require direct protein-level validation.

The reporter comparisons further show that fluorescent output in *Blastocystis* ST7-B is constrained by more than transgene expression alone, with each system exposing a different practical limitation. UnaG produced a clear signal in intact cells without oxygen-dependent chromophore maturation, which is a major advantage in an anaerobic protist (Kumagai et al., 2013; Kwon et al., 2020). However, UnaG fluorescence was compartmentalised despite the absence of an obvious targeting sequence, suggesting that signal distribution may be influenced by unconjugated BR availability or intracellular partitioning rather than reporter localization alone. This is plausible given the hydrophobic behaviour of unconjugated BR and its tendency to associate with lipid environments (Ostrow and Celic, 1984; Zucker et al., 1999; Bracken et al., 2024). Consistent with this, lipid-rich peripheral and intracellular structures have been reported in *Blastocystis* ST7-B, potentially providing favourable microenvironments for BR partitioning and contributing to the punctate UnaG fluorescence pattern (Liao et al., 2023). An alternative possibility is that the observed signal pattern reflects reporter sequestration or reporter-associated cellular stress. However, because UnaG-expressing lines were recovered, maintained under selection, and propagated through continued culture, toxicity remains a possible but untested explanation rather than a demonstrated mechanism. UnaG is therefore functional in *Blastocystis* ST7-B but appears less suitable for uniform whole-cell labelling or straightforward quantitative imaging across the cell. smURFP exposed a different constraint. Fluorescence was detectable only after saponin treatment, suggesting that limited chromophore access is likely the main barrier under the conditions tested (Rodriguez et al., 2016). This is consistent with the limited membrane permeability expected for BV and with the barrier properties attributed to the *Blastocystis* cell surface (Lightner et al., 1996; Bulmer et al., 2008; Yoshikawa and Hayakawa, 1996; Yason and Tan, 2018; Zheng and Gallot, 2020). In its current form, smURFP therefore appears better suited to permeabilised preparations than to straightforward live-cell imaging, although membrane-permeant bilins and newer monomeric variants may broaden its future utility (Hu et al., 2024; Maiti et al., 2023; Rodriguez et al., 2016). Among the reporters tested here, SNAP-tag® was the most adaptable in practice. Labelling worked in intact cells, signal scaled with substrate concentration, and the available chemistry provides access to a broad range of fluorophores and imaging modalities (Keppler et al., 2003; Gautier et al., 2008; Birke et al., 2022; Saimi and Chen, 2023; Porzberg et al., 2025). Although substrate-dependent labelling increases consumable cost, SNAP-tag® currently appears to offer the greatest practical versatility among the reporters evaluated here.

There are clear limitations to the present framework. The molecular maintenance state of the introduced constructs remains unresolved: episomal maintenance is a plausible working model, but genomic integration cannot be formally excluded. The constructs used here lack *Blastocystis* ST7-B homology arms, and comparative genomic analyses suggest that *Blastocystis* lacks canonical non-homologous end-joining components (Gentekaki et al., 2017), making targeted integration by standard repair routes unlikely but not impossible. Direct assays such as outward-facing PCR, plasmid rescue followed by full plasmid sequencing, FISH, or selection-withdrawal experiments will be required to distinguish episomal persistence from integration. Copy number, segregation, and long-term expression stability remain unresolved, and these variables are likely to contribute to clone-to-clone heterogeneity, particularly where more quantitative readouts are required. The toolkit also remains limited to transgene expression and reporter validation, with targeted integration, conditional control, and direct loss-of-function approaches still to be established.

## Conclusions

Taken together, this work establishes a practical framework for stable transgenesis under antibiotic selection in *Blastocystis* ST7-B, substantially expanding the genetic tools available for this organism. By integrating endogenous promoter and terminator discovery, DNA delivery, selection, clonal recovery, and reporter validation into a single pipeline, we provide a flexible foundation for routine transgene-based studies. Our results show that *Blastocystis* ST7-B can be genetically modified and that stable, colony-derived transgenic lines can be generated, opening the way to more direct molecular analysis of its cell biology. More broadly, this strategy offers a practical blueprint for developing genetic tools in other less understood microbial eukaryotes where genetic access remains limited.

## Acknowledgements

MRT was supported by Norwegian Research Council grant 301170 to MvdG. MRT is grateful for COST Action CA21105 (*Blastocystis* under One Health) for part-sponsoring his visit to KSWT’s laboratory. We thank Ms. Geok Choo Ng and Dr. Steven Santino Leonardi for generously sharing their wet-lab techniques for culturing Blastocystis. We thank Dr. Yohannes Seyoum for helping with the statistical analyses. We also thank Luz Aurora Martinez-Contreras, Daniel Simon Nagel, and Ayanara Mae Sadi Bukneberg for their help in routine culturing and maintenance of *Blastocystis* cultures.

## Author contributions

Conceptualisation and design of the research: MRT and MvdG

Critical advice: MvdG and KSWT

Experimental work and data analyses: MRT

Writing and editing: MRT, MvdG, and KSWT

All authors read and revised the paper.

## Supplemental Information

**Supplemental Data 1.** Gene sequences of reporter genes, selection markers, Kozak, and P2A linker

>JQ437370.1 NanoLuc luciferase

ATGGTCTTCACACTCGAAGATTTCGTTGGGGACTGGCGACAGACAGCCGGCTACAACCTGGACCAAGTCCTTGAACAGGGAGGTGTGTCCAGTTTGTTTCAGAATCTCGGGGTGTCCGTAACTCCGATCCAAAGGATTGTCCTGAGCGGTGAAAATGGGCTGAAGATCGACATCCATGTCATCATCCCGTATGAAGGTCTGAGCGGCGACCAAATGGGCCAGATCGAAAAAATTTTTAAGGTGGTGTACCCTGTGGATGATCATCACTTTAAGGTGATCCTGCACTATGGCACACTGGTAATCGACGGGGTTACGCCGAACATGATCGACTATTTCGGACGGCCGTATGAAGGCATCGCCGTGTTCGACGGCAAAAAGATCACTGTAACAGGGACCCTGTGGAACGGCAACAAAATTATCGACGAGCGCCTGATCAACCCCGACGGCTCCCTGCTGTTCCGAGTAACCATCAACGGAGTGACCGGCTGGCGGCTGTGCGAACGCATTCTGGCGTAA

>smURFP *Blastocystis* ST7-B codon-optimized

ATGGCTAAGACGTCGGAGCAGCGCGTGAACATTGCGACGCTGCTGACGGAAAACAAGAAAAAGATCGTGGATAAGGCGTCGCAGGATCTGTGGCGACGCCATCCGGATCTGATTGCGCCTGGCGGAATCGCGTTCTCGCAGCGCGATCGAGCGCTGTGCCTGCGCGATTACGGATGGTTCCTGCACCTGATCACGTTCTGCCTGCTGGCTGGCGATAAGGGACCGATCGAGTCGATTGGACTGATCTCGATCCGCGAGATGTATAACTCGCTGGGAGTGCCGGTGCCTGCTATGATGGAATCGATCCGCTGCCTGAAAGAGGCGTCGCTGTCGCTGCTGGATGAGGAAGATGCGAACGAGACTGCGCCTTACTTCGATTACATCATCAAGGCTATGTCGTAA

>SNAP-tag *Blastocystis* ST7-B codon-optimized

ATGGATAAGGATTGCGAGATGAAGCGCACGACGCTGGATTCGCCTCTGGGAAAGCTGGAACTGTCGGGATGCGAGCAGGGACTGCACCGCATCATCTTTCTCGGAAAGGGAACGTCGGCTGCGGATGCGGTGGAAGTGCCTGCGCCTGCTGCTGTGCTCGGAGGACCTGAGCCTCTGATGCAGGCGACGGCGTGGCTGAACGCGTACTTCCATCAGCCTGAGGCGATCGAGGAATTCCCGGTGCCTGCTCTGCACCATCCGGTGTTCCAGCAAGAGTCGTTCACGCGACAGGTGCTGTGGAAGCTGCTGAAGGTGGTGAAGTTCGGAGAGGTGATCTCGTACTCGCACCTGGCTGCGCTGGCTGGAAACCCTGCGGCTACGGCTGCGGTGAAAACGGCTCTGTCGGGAAACCCGGTGCCGATCCTGATTCCGTGCCACCGCGTGGTGCAGGGCGATCTGGATGTTGGAGGATACGAAGGCGGACTGGCGGTGAAAGAGTGGCTGCTGGCGCACGAGGGACACCGCCTCGGAAAGCCTGGACTGGGATAA

>UnaG *Blastocystis* ST7-B codon-optimized

ATGGTCGAGAAGTTCGTTGGAACGTGGAAGATCGCGGATTCGCACAACTTCGGAGAGTACCTGAAGGCGATCGGAGCGCCTAAAGAACTGTCTGATGGCGGAGATGCTACGACGCCGACGCTGTACATTTCGCAAAAGGATGGCGATAAGATGACCGTGAAGATCGAGAACGGACCGCCGACCTTCCTGGATACGCAAGTGAAGTTCAAGCTGGGAGAAGAGTTCGATGAGTTCCCGTCGGATCGCCGAAAGGGCGTGAAGTCGGTGGTTAACCTCGTGGGAGAGAAGCTGGTGTACGTGCAGAAGTGGGACGGAAAAGAAACGACCTACGTGCGCGAGATCAAGGATGGAAAGCTGGTGGTTACGCTGACGATGGGAGATGTGGTGGCTGTGCGATCGTATCGCCGAGCGACTGAATAA

>E. coli dhfr *Blastocystis* ST7-B codon-optimized

ATGGAAAACGCTATGCCGTGGAACCTGCCTGCGGATCTGGCGTGGTTCAAGCGCAACACGCTGAACAAGCCGGTGATCATGGGACGCCACACGTGGGAGTCGATCGGACGCCCTCTGCCTGGACGCAAGAACATCATCCTGTCGTCGCAGCCTGGAACGGATGACCGCGTGACGTGGGTGAAGTCGGTGGATGAGGCGATTGCGGCTTGCGGAGATGTGCCCGAGATCATGGTGATCGGAGGCGGACGCGTGTACGAGCAGTTCCTGCCGAAGGCGCAGAAGCTGTACCTGACGCACATCGATGCGGAGGTGGAAGGCGATACGCACTTCCCGGATTACGAGCCGGATGATTGGGAGAGCGTGTTCTCGGAGTTCCACGATGCGGATGCGCAGAACTCGCACTCGTACTGCTTCGAGATCCTGGAACGCC GATAA

>human DHFR *Blastocystis* ST7-B codon-optimized

ATGCACGGATCGCTGAACTGCATCGTGGCGGTGTCGCAGAACATGGGAATCGGAAAGAACGGCGATTACCCGTGGCCTCCGCTGCGCAACGAGTTCCGCTACTTCCAGCGCATGACGACGACGTCGAGCGTGGAAGGCAAGCAGAACCTGGTGATCATGGGAAAGAAAACGTGGTTCTCGATCCCCGAGAAGAATCGCCCTCTGAAGGGACGCATCAACCTGGTGCTGTCGCGAGAGCTGAAAGAACCGCCTCAGGGTGCGCACTTCCTGTCGCGCTCGCTGGATGATGCGCTGAAGCTGACGGAACAGCCCGAGCTGGCGAACAAGGTGGATATGGTGTGGATCGTTGGAGGATCGTCGGTGTACAAAGAAGCTATGAACCATCCGGGACACCTGAAGCTGTTCGTGACGCGCATCATGCAGGATTTCGAGTCGGATACGTTCTTCCCCGAGATCGACCTGGAAAAGTACAAGCTGCTGCCTGAGTACCCTGGCGTGCTGTCGGATGTGCAAGAGGAAAAGGGAATCAAGTACAAGTTCGAGGTGTACGAGAAGAACGACTAA

>pac *Blastocystis* ST7-B codon-optimized

ATGACCGAGTACAAGCCTACTGTGCGACTGGCGACGCGAGATGATGTTCCTAGAGCTGTGCGAACGCTGGCGGCTGCGTTTGCTGATTATCCTGCTACGCGCCACACGGTGGATCCTGATCGCCATATTGAACGCGTGACGGAACTGCAAGAGCTGTTCCTGA CGCGCGTGGGACTCGATATCGGAAAAGTGTGGGTTGCCGATGATGGCGCTGCTGTGGCTGTGTGGACTACGCCTGAATCTGTGGAAGCGGGTGCTGTGTTCGCGGAGATTGGACCTAGAATGGCGGAACTGTCTGGATCTAGACTGGCGGCGCAGCAGCAGATGGAAGGACTTTTGGCGCCGCACCGACCTAAAGAACCTGCTTGGTTTCTGGCGACCGTGGGAGTGTCTCCGGATCACCAAGGCAAAGGATTGGGATCTGCTGTGGTGCTGCCTGGCGTTGAAGCTGCTGAACGAGCTGGTGTTCCGGCTTTCTTGGAAACGTCGGCGCCTCGCAACCTGCCTTTCTATGAACGCCTGGGATTCACCGTGACGGCGGATGTGGAAGTTCCTGAAGGACCTCGAACGTGGTGCATGACGCGCAAGCCTGGTGC TTAA

>P2A sequence *Blastocystis* ST7-B codon-optimized

GGCAGCGGAGCGACGAACTTCTCGCTGCTGAAGCAGGCTGGCGACGTGGAAGAGAACCCTGGACCG

---

**Supplemental Table 1.**
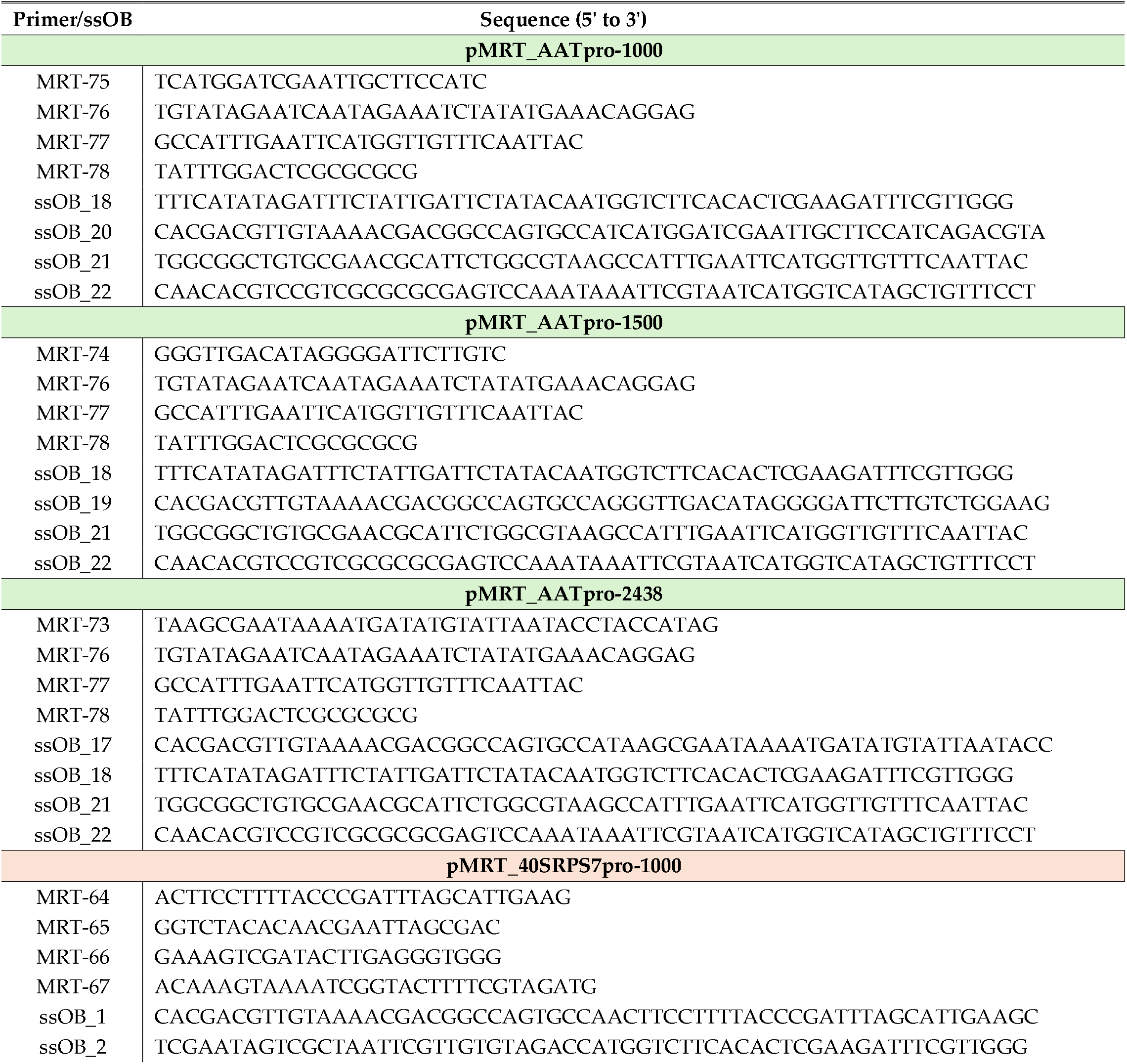
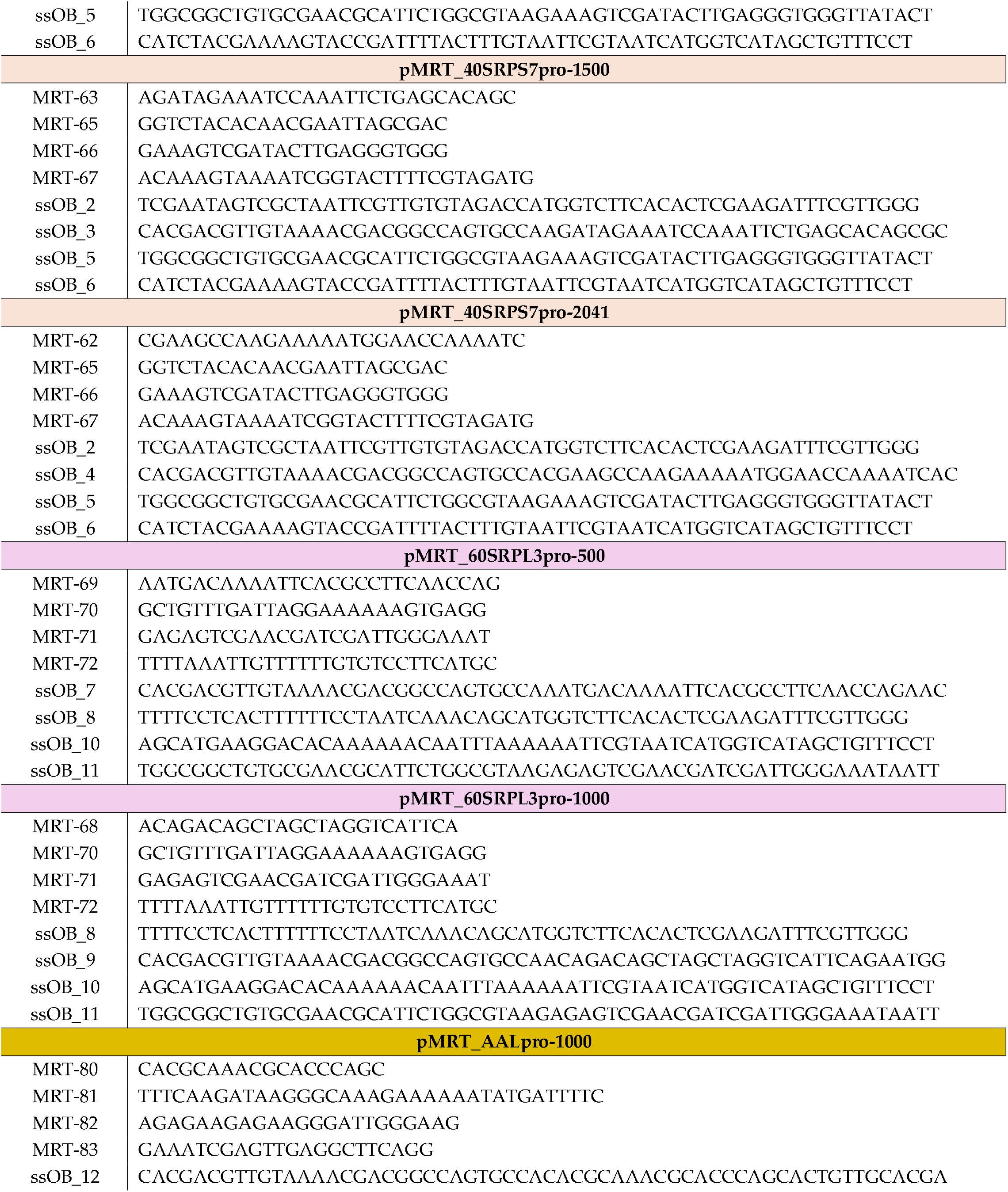
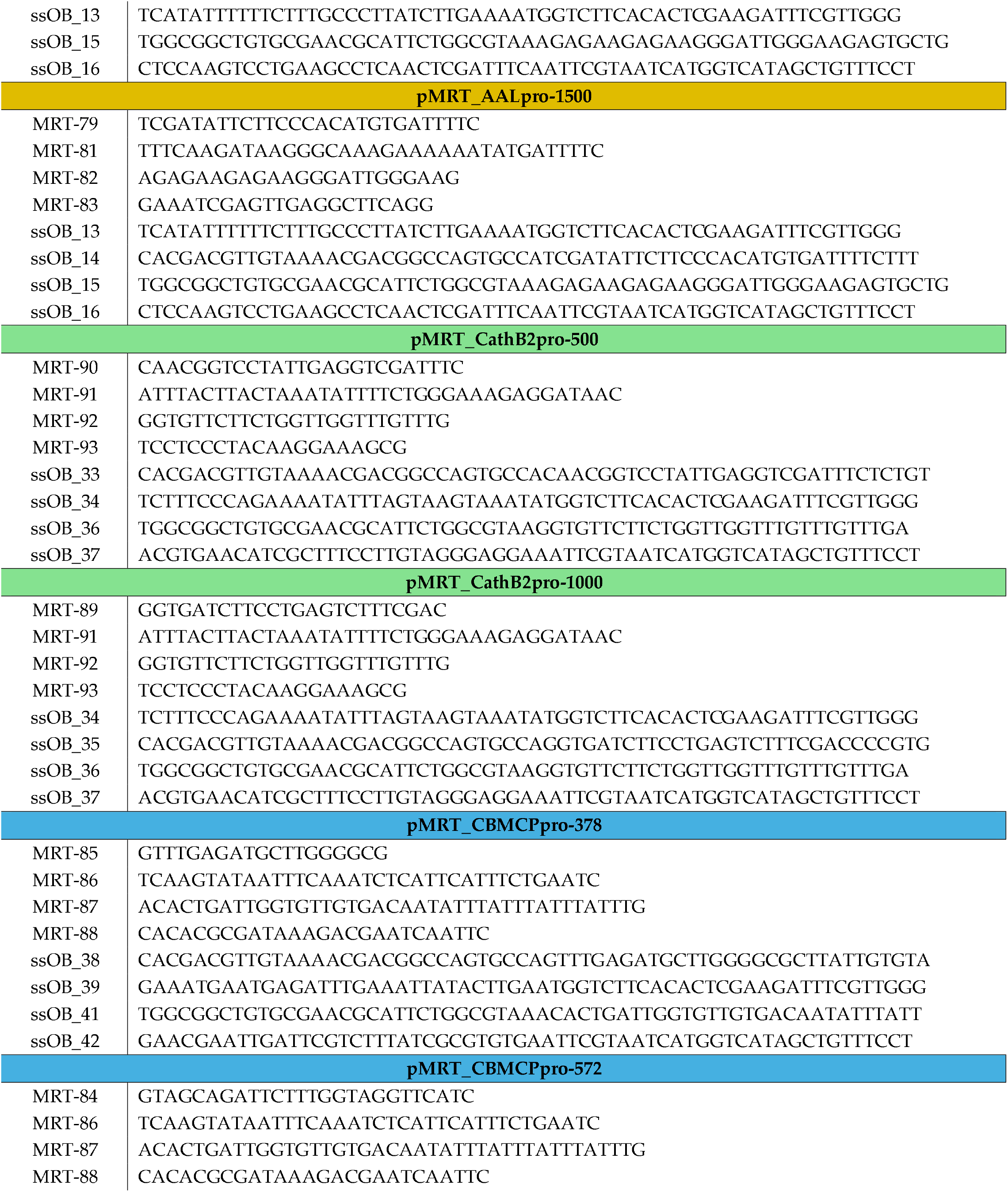
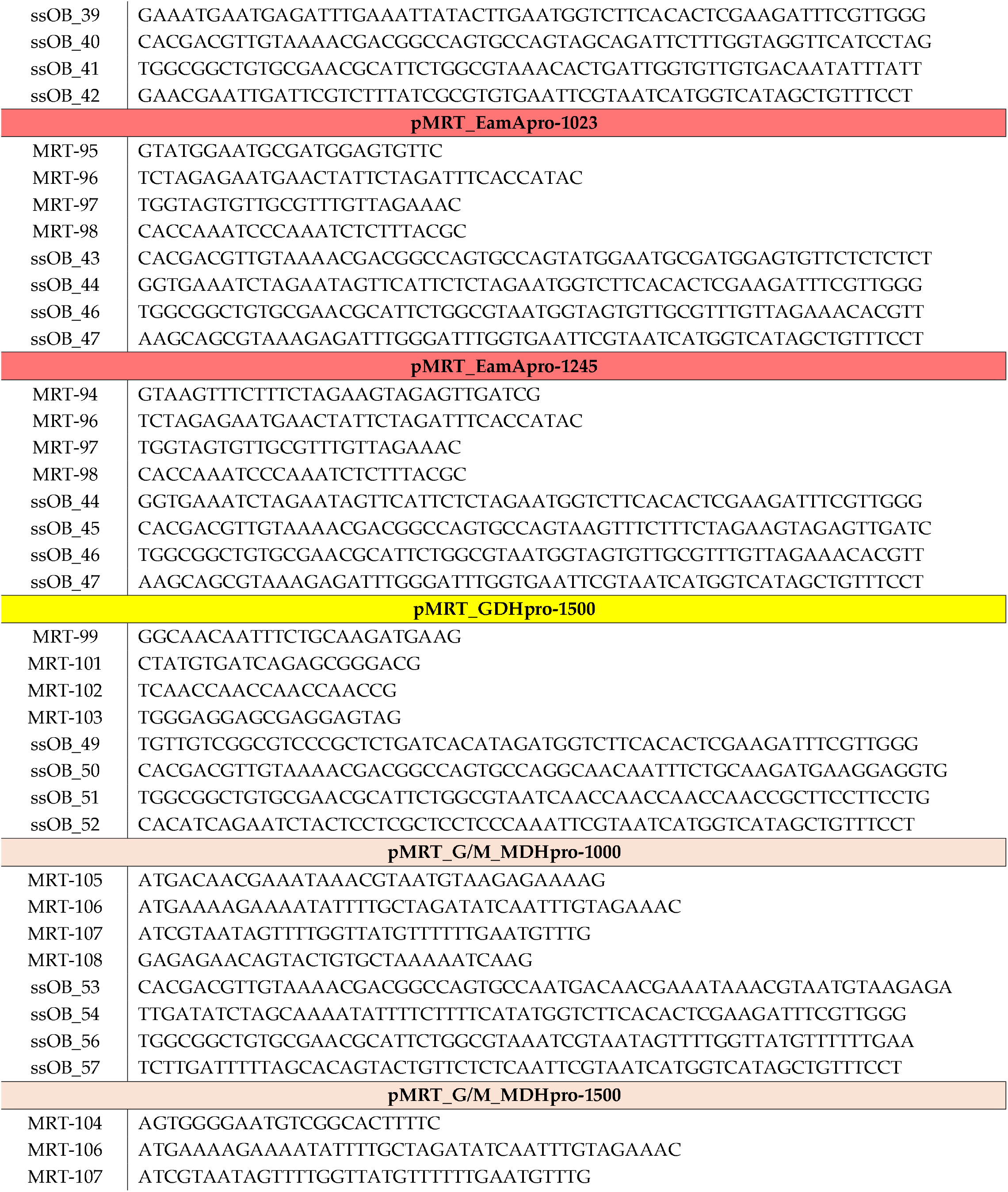
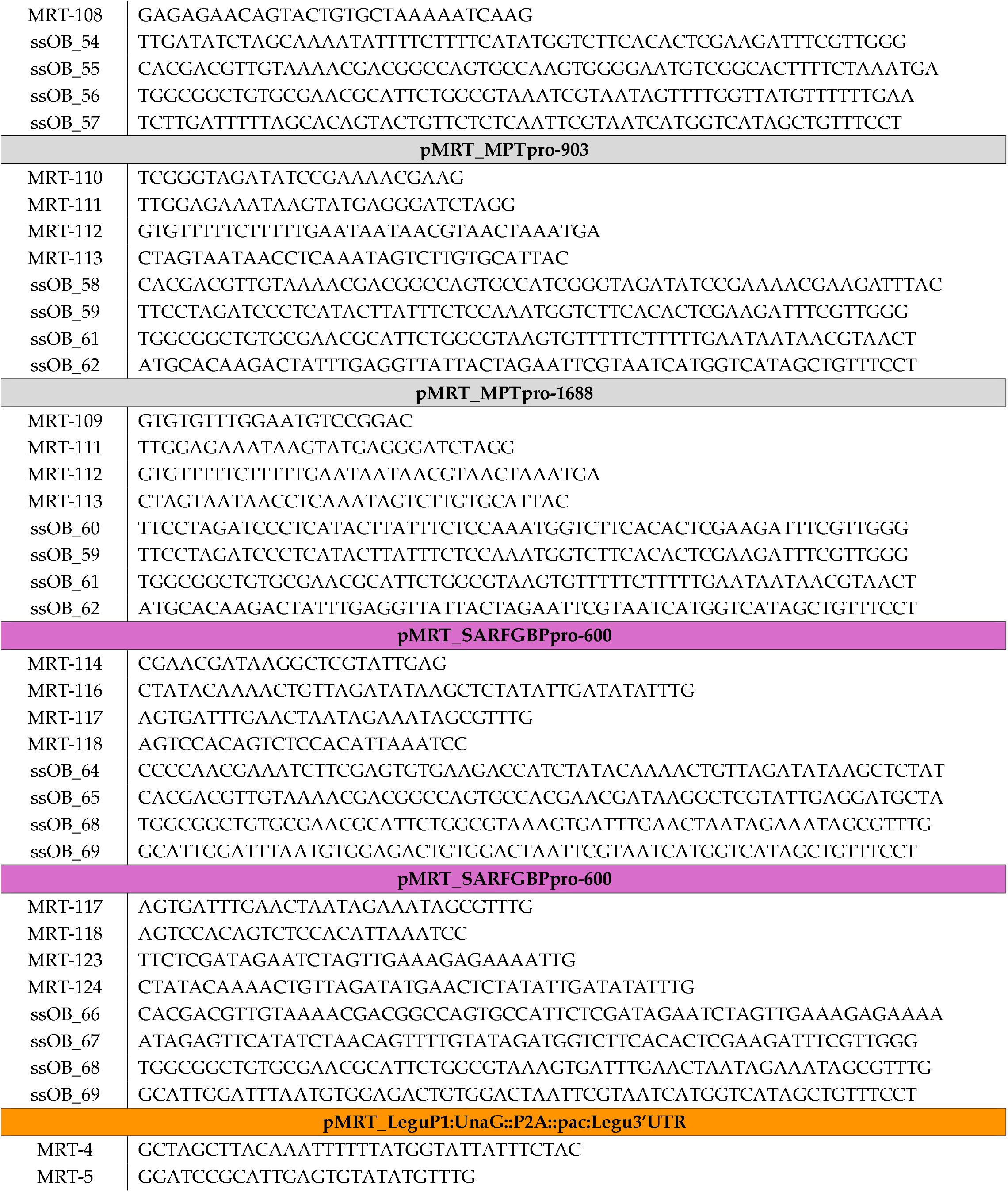
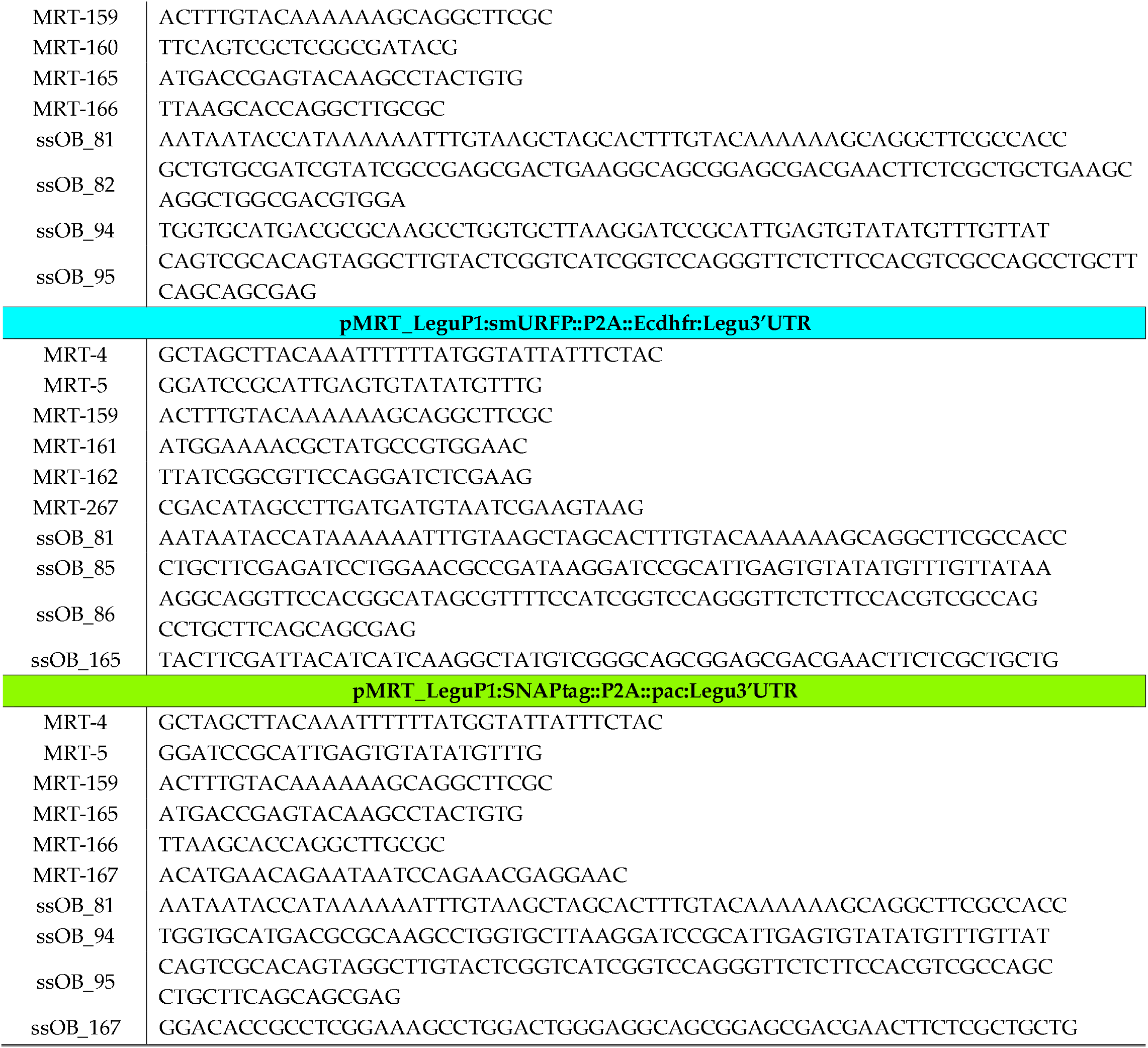
Oligonucleotide sets used for expression vector cloning. In cases where only the promoter fragment was changed, most oligonucleotides were reused; however, complete sets are shown here for clarity and completeness. Primers used for PCR amplification of DNA fragments are designated MRT-xxx, whereas single-stranded oligonucleotide bridges used for DNA assembly are termed ssOB-yyy. Oligonucleotides are grouped by plasmid, named pMRT-zzz.

